# Ancestry-specific performance of variant effect predictors in clinical variant classification

**DOI:** 10.64898/2026.02.14.705914

**Authors:** Rachel Hoffing, Daniel Zeiberg, Sarah L. Stenton, Matthew Mort, David N. Cooper, Matthew W. Hahn, Anne O’Donnell-Luria, Lucas D. Ward, Predrag Radivojac

## Abstract

Predicting the effects of genetic variants and assessing prediction performance are key computational tasks in genomic medicine. It has been shown that well-calibrated variant effect predictors can be reliably used as evidence towards establishing pathogenicity (or benignity) of missense variants, thereby rendering these variants suitable for use in (or exclusion from) the genetic diagnosis of rare Mendelian conditions. However, most predictors have been trained or calibrated on data that may not be sufficiently representative to lead to similar performance across all genetic ancestries. This raises questions about the responsible deployment of these tools to improve human health. To better understand the utility of computational predictors, we set out to assess their ancestry-specific performance in terms of accuracy and evidence strength according to the ACMG/AMP guidelines. First, we determined that the expected count of rare variants in an individual’s genome and the allele frequency distribution of these variants are the key confounders when evaluating a predictor’s performance across different genetic ancestries. Second, we found that a predictor’s accuracy itself inversely correlates with the allele frequency of the rare variant. After stratifying according to allele frequency, we show that established methods for predicting the pathogenicity of missense variants have comparable performance levels across major ancestry groups. Our results therefore support the wide deployment of such models in the context of genetic diagnosis and related applications.

## Introduction

Variant effect predictors provide a powerful *in silico* approach to assess the impact of rare missense variation on the function of gene products [18, 24, 27]. Owing to their scalability and high accuracy [50, 38], carefully calibrated tools allow for a genome-wide application [34, 45, 3] and can facilitate the clinical classification of variants in disease-associated genes as “pathogenic” or “benign” [39, 52]. Scalability and high accuracy, however, are often insufficient for the real-world deployment of machine learning models [31]. To be broadly regarded as reliable for use in genomic medicine, it is also essential that the performance of these tools does not vary based on an individual’s genetic ancestry.

The importance of this type of evaluation has been apparent for methods such as polygenic risk scores (PRS), which aim to predict phenotypic outcomes in complex disease based on a sum of estimated effects of common risk alleles [54]. PRS are built on genome-wide association studies, which have been conducted primarily among individuals of European genetic ancestry [37, 42, 16] and often translate poorly to non-European genomes [29, 12, 28]. The same types of transferability concerns remain largely unaddressed in the field of rare genetic disease, where hundreds of models have been published to date [27] and several have been recommended for clinical use [34, 3].

Few studies have investigated ancestry-specific variant effect predictor evaluation. Parthak et al. [33] found ancestry-specific differences in score distributions for a subset of models directly trained or evaluated on variants labeled as “pathogenic” and “benign” in major databases such as ClinVar [26], Human Gene Mutation Database (HGMD) [44] and UniProt [2]. By contrast, tools trained on data without population-level information, such as functional data from MaveDB [14], showed smaller ancestry-specific discrepancies, suggesting problems with the use of tools trained on population-dependent labeled data. Dawood et al. [9] reported similar concerns regarding computational predictors, as well as higher frequencies of variants of uncertain significance in individuals of non-European ancestry, higher rates of benign variants, but lower rates of pathogenic variants. They argue that disparities in variant classification can be reduced by expanding deep mutational scanning experiments. Most recently, De Paolis Kaluza et al. [10] developed a formal semi-supervised procedure that improves the estimation of fairness metrics for machine learning models when labeled data is biased. In contrast to previous studies, their results on a single variant effect predictor trained on population data show only minor performance discrepancies between broad ancestry groups. Although this bias-correction approach does not require the knowledge of ancestry for labeled variants, it does require the availability of feature vectors (variant embeddings) for both labeled and unlabeled variants, which limits its use to a very small group of existing models (e.g., MutPred2 [35], ESM1b [4]).

In this study, we relied on the data from 425,978 exome-sequenced participants from the UK Biobank [1], along with variant data from gnomAD [22], ClinVar [26], and HGMD [44] to provide a comprehensive investigation of the ancestry-specific performance of several prominent models for predicting the effects of missense variation. We first determined that the number of rare missense variants in an individual’s genome and the allele frequency distribution among those variants are key confounders in the analysis of ancestry-specific performance. We then noted large differences in the classification accuracy of the variant effect predictors currently recommended for clinical use [34, 3] when applied to rare variants in different ranges of (estimated) allele frequency. Finally, we provide evidence that, when the analysis is stratified according to allele frequency, the evaluated tools were broadly fair, thus supporting their deployment for clinical variant classification across diverse populations.

## Results

### Heterozygous rare missense variant sites per person

To assess the performance of variant effect prediction tools across individuals of different ancestries, we first examined the overall extent of rare genetic variation in Admixed American, African, Central/South Asian, East Asian, European, and Middle Eastern populations. To achieve this, we compiled rare variants, defined here as those with an alternate allele frequency (AF) below 1% across all ancestries in gnomAD, together with their evolutionary conservation scores at the corresponding genomic positions. We then computed the degree of heterozygosity of missense variants within a given gene set, with synonymous variants in the same genes acting as control for neutral variation. In addition, we computed the average AF and evolutionary conservation over those variants for each individual. Since our primary interest is clinical variant interpretation in the context of rare disease, our main analyses focus on a curated set of genes (*n* = 4,780) enriched for disease association, as described in Methods, while results for all protein-coding genes are provided in Supplementary Table S1.

Figure 1 shows the extent of heterozygous synonymous (Figure 1a,b) and missense (Figure 1a,d) variation in the set of genes enriched for disease association. Because we have the largest number of European-ancestry individuals, we have the largest number of variants from this group (Figure 1a). Nevertheless, we observe the highest degree of heterozygosity among individuals of African genetic ancestry (missense: 161±17 [mean ± standard deviation]; synonymous: 131±16) for both missense and synonymous variants, followed by the Central/South Asian (missense: 125 ± 13; synonymous: 84±11), East Asian (missense: 119±20; synonymous: 79±14), Middle Eastern (missense: 113±33; synonymous: 80 ± 28), European (missense: 94 ± 10; synonymous: 59 ± 8), and Admixed American (missense: 90 ± 16; synonymous: 62 ± 12) ancestries. These results are consistent with previous investigations [49, 48].

**Figure 1.**
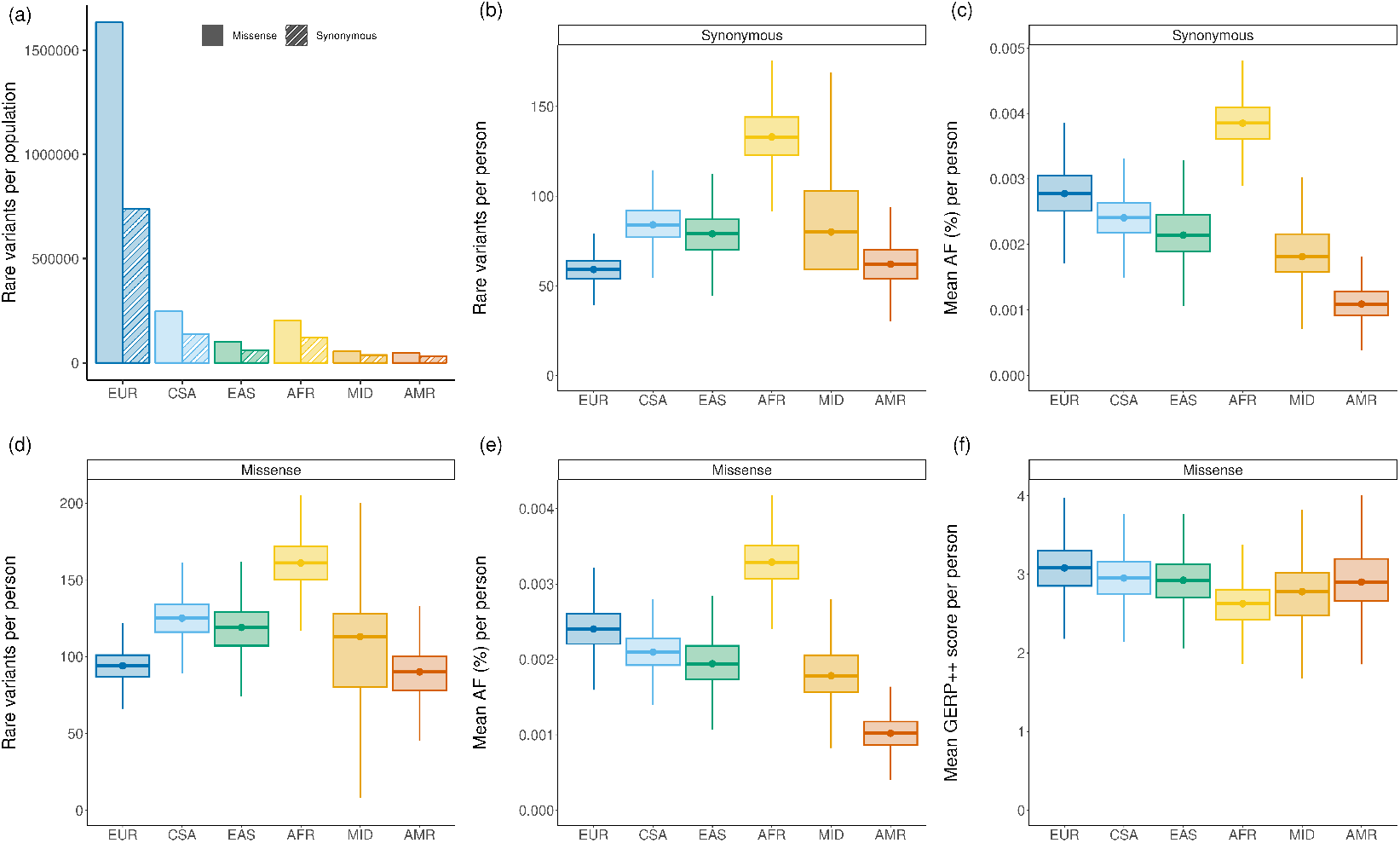
Each boxplot represents rare (AF < 1%) variants in genes enriched for disease-association, stratified by genetic ancestry (Admixed American = AMR, African = AFR, Central/South Asian = CSA, East Asian = EAS, European = EUR, and Middle Eastern = MID). Boxplots depict the median value as the center, first and third quartiles as box boundaries, and whiskers extending 1.5-times the inter-quartile range with outliers excluded, and values bounded at 0 for total variants and AF. (a) Total rare missense and synonymous variants by genetic ancestry. (b) Rare heterozygous synonymous variants per person. (c) AF of rare heterozygous synonymous variants per person. (d) Rare heterozygous missense variants per person. (e) Mean AF of rare heterozygous synonymous variants per person. (f) Mean GERP++ rank score of rare heterozygous missense variants per person (higher score indicates higher evolutionary conservation).

We also looked at the estimated AF for these variants (Figure 1c,e) and the evolutionary constraint at the missense variant positions (Figure 1f), as computed from the multiple sequence alignments incorporating the reference human genome along with those of multiple other species. Although the accuracy of estimated AF for each ancestry is differentially influenced by the population sample size in gnomAD, Figure 1c and Figure 1e demonstrate considerable differences in the rare variant AF distribution across ancestral groups. Synonymous and missense variants follow a similar pattern, with missense variants showing slightly lower average AF than synonymous. Finally, Figure 1f shows that the mean GERP++ [8] scores at the positions of rare missense variation in the same group of genes are inversely correlated with the number of variants in each group, indicating the highest degree of conservation in individuals of European ancestry, followed by Central/South Asian, Admixed American, East Asian, Middle Eastern and African ancestries. The results for all genes and those using the PhyloP [36] conservation scores are provided in Supplementary Table S1.

### Ancestry-specific clinical evidence yield for rare missense variants

Extending the analysis of Stenton et al. [45] to genetic ancestries, we counted rare missense variants in a genome stratified according to their evidence strength within the American College of Medical Genetics and Genomics and Association for Molecular Pathology (ACMG/AMP) sequence variant classification guidelines [39]. Three computational predictors, REVEL [21], MutPred2 [35] and AlphaMissense [7], were used to generate predictions on all individuals, and the variants were then categorized into PP3/BP4 evidence levels according to the recent ClinGen recommendations [34, 3]. PP3/BP4 evidence codes correspond to the evidence direction (pathogenic vs. benign) and strength (both qualitatively reported as supporting, moderate, strong and quantitatively reported as ±1 to ±4 points) recommended for computational predictions. Computational evidence is then combined with other types of evidence (e.g., population, functional) to label variants observed in rare-disease patients as pathogenic, likely pathogenic, likely benign, and benign [39]. Pathogenic and likely pathogenic variants are considered diagnostic and variants labeled benign or likely benign are excluded as a genetic diagnosis. The remaining variants observed in patients with rare disease are labeled as variants of uncertain significance (VUS).

Figure 2a shows the distribution of the number of variants within genes enriched for disease association averaged over the three models and stratified according to the quantitative clinical evidence levels and ancestral groups. We observe that the PP3 categories are consistent across the groups, with increasing variability as the evidence strength decreases. Depending on the ancestry group, we find an average of 1.6-1.8 variants with PP3-strong (+4), 4.8-5.9 with PP3-moderate (+2, +3) and 4.2-5.7 with PP3-supporting (+1) evidence per individual. This implies that the discrepancy in evidence strength yield across ancestries increases with the increasing fraction of false positive predictions in each evidence strength category (largest false positive rate for PP3-supporting, smallest for PP3-strong [34]). Consistent with this pattern, BP4 evidence exhibits larger variability that mirrors the overall burden of rare missense variation among ancestries. Depending on the ancestry group, we find an average of 18-33 variants with BP4-supporting and 32-61 with BP4-moderate (*−*2, *−*3) evidence per individual; see Figure 2a. We do not show the BP4-strong category, as a very small fraction of variants can be assigned this evidence strength and some of the predictors do not even reach this evidence category [3]. The results follow the same trend over all genes and are similar for the remaining predictors. Complete results for all individual predictors are given in Supplementary Table S1.

**Figure 2.**
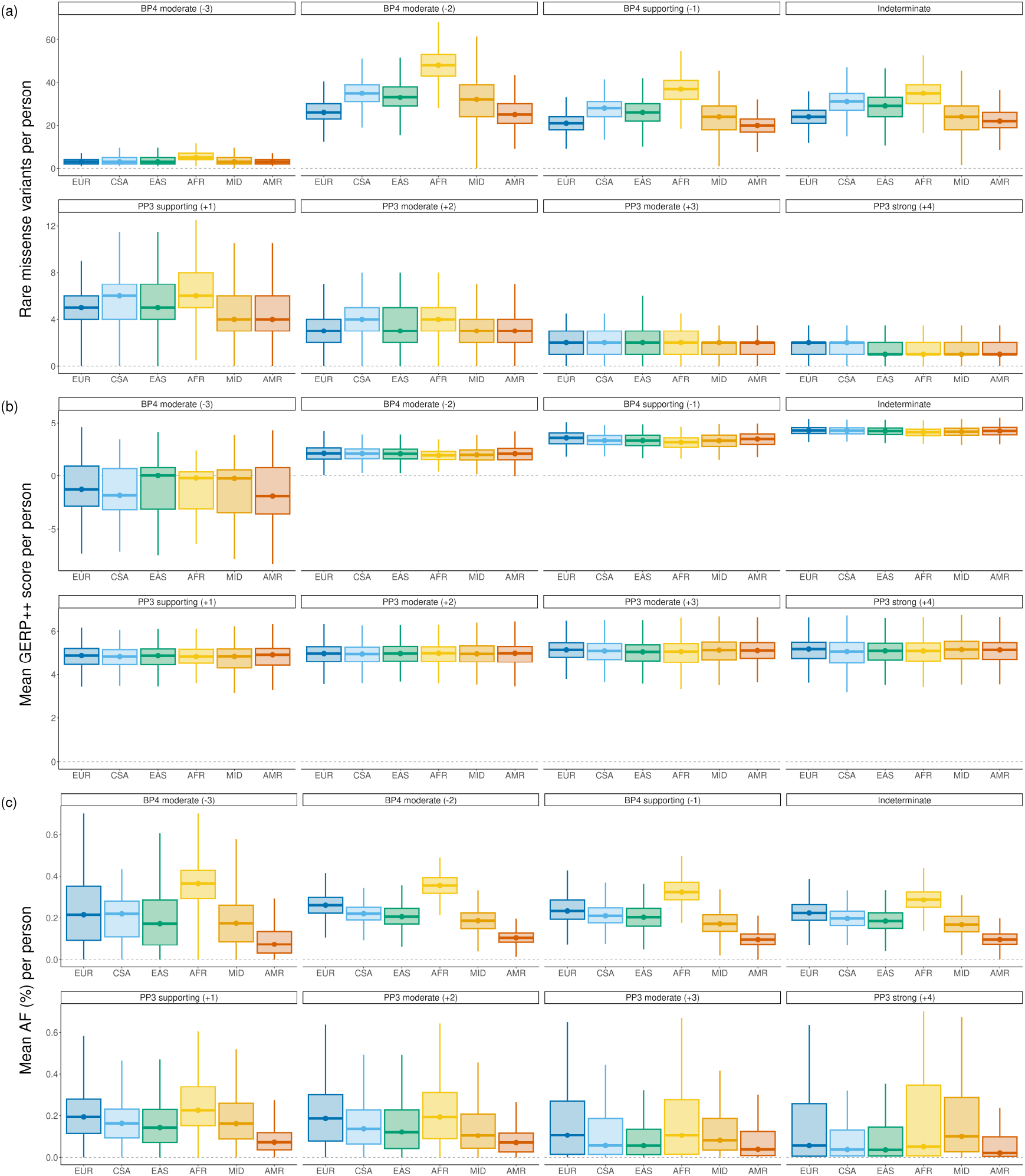
Each boxplot represents the averaged value across REVEL, MutPred2, and AlphaMissense predictions for rare (AF < 1%) missense variants in disease genes, stratified by ACMG/AMP guidelines for pathogenic and benign evidence and genetic ancestry (Admixed American = AMR, African = AFR, Central/South Asian = CSA, East Asian = EAS, European = EUR, and Middle Eastern = MID). Boxplots depict the median value as the center, first and third quartiles as box boundaries, and whiskers extending 1.5-times the inter-quartile range with outliers excluded, and values bounded at 0 for total variants and AF. Results show (a) Rare heterozygous missense variants per person (b) Median GERP++ rank score of rare heterozygous missense variants per person (c) Median allele frequency (%) of rare heterozygous missense variants per person.

We next sought to explain the reasons why PP3 and BP4 evidence levels do not both show the same pattern; that is, why is PP3 evidence relatively constant across the groups, while BP4 evidence is variable. Since evolutionary information provides the main signal for variant effect predictors [5], we analyzed the distribution of per-individual mean GERP++ scores across evidence levels and genetic ancestries. Figure 2b shows that the variants predicted within PP3 evidence classes are similarly consistent in their evolutionary conservation within evidence level classes. On the other hand, BP4 categories show increased variability and trends consistent with the distribution of the number of variants in each population. Importantly, conservation levels steadily increase across all predictors with the evidence levels increasing from *−*3 to +4. Similar results can be observed using estimated AF (Figure 2c). However, we caution that these results have differentially accurate AF estimates across populations due to uneven ancestry representation in population databases—over two orders of magnitude size difference between the smallest and the largest [22].

Overall, these results demonstrate that variant effect prediction is largely driven by evolutionary conservation and suggest that evolutionary conservation, rather than ancestry, is the main driver in variant effect prediction. Interestingly, this is true irrespective of how the variant effect predictor was trained. While models such as REVEL and MutPred2 were trained in a supervised manner between pathogenic and benign (or unlabeled) variants, AlphaMissense did not use such supervision, yet it effectively identifies evolutionary signals. A larger trend that a number of highly diverse computational models correlate more among themselves than either of them does with low- or high-throughput experimental data has been previously reported [50].

### Prediction performance on ClinVar and HGMD pathogenic and benign variants

To establish the predictor performance across genetic ancestries, we next collected known pathogenic and benign variants from ClinVar [26] and HGMD [44]. All rare variants with a pathogenic, likely pathogenic, benign, or likely benign status in ClinVar with at least a one-star rating, as well as disease-causing mutation (DM) variants in HGMD, were included in the analysis. Benign variants were collected only from genes enriched for disease association. From these datasets, we assigned ancestry to each variant based on its presence in gnomAD and removed variants not observed in any population (a variant was included in the data for every ancestry group in which it was observed in gnomAD). We then evaluated how well each predictor distinguished pathogenic from benign variants within each group by calculating the area under the Receiver Operating Characteristic curve (AUC); see Figure 3 and more detailed results in Supplementary Table S2.

**Figure 3.**
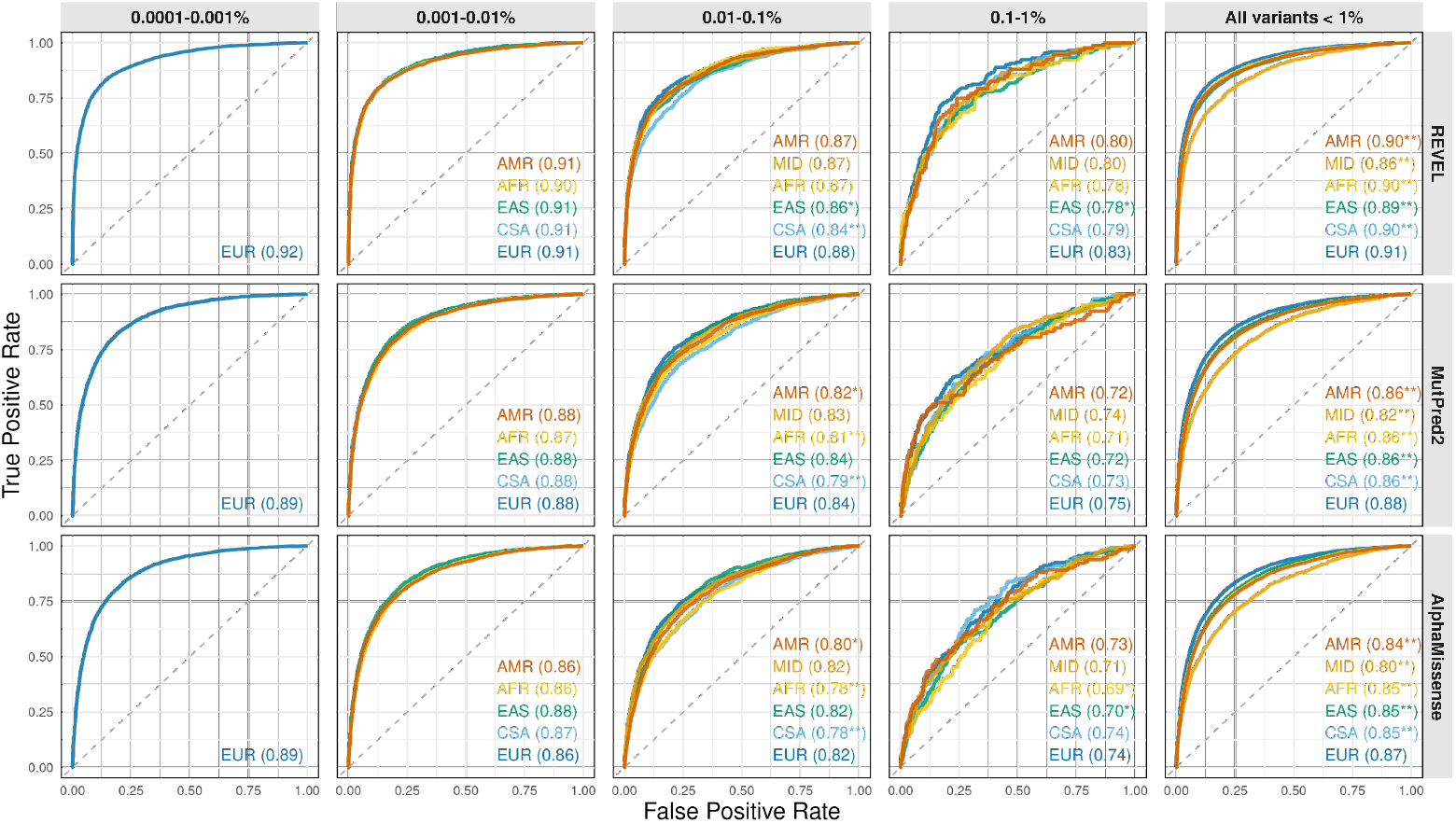
Performance of REVEL, MutPred2 and AlphaMissense against labeled ClinVar and HGMD variants, binned by allele frequency and stratified by genetic ancestry. Bootstrapping was conducted over 1,000 iterations to compute a P-value. Star annotation indicate P-value significance at *P* < 0.05 (*) and Bonferroni-adjusted P-value per group at *P* < 0.05/19 (**).

We carried out this analysis overall as well as binned by allele frequency. In the overall analysis, there were relatively more pathogenic variants in the European sample. This is because a substantially larger sample size of European-ancestry individuals in both gnomAD and the UK Biobank enriches the analysis for very rare variants, which are more likely to be deleterious (Figure 2) and more likely to be accurately predicted as such (Figure 3). To determine if the performance of these models across ancestral groups showed statistical differences, we carried out bootstrapping over 1,000 iterations [13], stratifying by both ancestry and allele frequency. We determined empirical P-values by calculating the number of instances in which the non-European AUC exceeded that of the European result, divided by 1,000 bootstrapped iterations.

The results of the analysis binned by allele frequency suggest similar performance of each tool across different ancestries, with only 5 out of 42 European-to-non-European comparisons showing statistical significance after Bonferroni correction. On the other hand, when all variants were combined (Figure 3, last column), virtually all 15 comparisons between European and non-European ancestries show statistical significance, suggesting Simpson’s reversal due to confounding by the allele frequency effects on accuracy and differential group sizes. The absence of ancestry-specific effects after accounting for allele frequency indicates that these predictors can be applied across human populations without introducing bias. It also implies that comparisons of prediction performance between groups should only be made after conditioning on allele frequency.

## Discussion

When setting the group fairness expectations for computational variant effect predictors, it is imperative to carefully consider the differences between ancestral groups in terms of the expected number of variants per individual as well as the allele frequency distribution of these variants. Would a fair predictor performance suggest the same performance accuracy over all variants observed in these ancestries? Should all ancestry groups show similar rates or similar numbers of predicted pathogenic variants per individual, or should there be a more nuanced expectation? What should be the expectation for benign variation? Because the major fairness conditions we can impose on computational tools are difficult to satisfy, and fairness metrics cannot be optimized simultaneously [25, 31], we must at least make an attempt at their accurate estimation before making a judgment. However, this is challenging due to both the potential biases in the collection of labeled data and the absence of ancestry information in ClinVar and HGMD. Although some techniques for bias correction in the estimation of fairness metrics have been recently developed [10], they can be applied to only two out of seven predictors currently recommended for clinical use [34, 3].

We therefore sought alternative ways to understand the ancestry-specific performance of variant effect predictors for the purposes of clinical variant classification. We first determined the baseline per-genome load of rare missense variation in the UK Biobank across six genetic-ancestry samples and identified considerable differences in allele frequency distributions among those groups of variants. In line with earlier studies [49, 48], we found that the individuals of European genetic ancestry had a relatively low number of rare missense variants and the positions of these variants exhibit the highest degree of evolutionary conservation among all ancestries (Figure 1). On the other hand, individuals of African genetic ancestry had a comparably high amount of variation to other ancestries and the positions of rare variants in this population exhibit the lowest degree of average evolutionary conservation (Figure 1).

We next evaluated the evidence yield per individual as specified by ACMG/AMP guidelines, stratifying by population. We found that the assigned benign evidence closely followed the overall variant counts within ancestries (Figure 2a, BP4 panels). Among variants with assigned pathogenic evidence, we found that all ancestral groups have similar numbers of variants per genome (Figure 2a, PP3 panels). At the most restrictive evidence threshold, each genome contains approximately 1.6-1.8 PP3-strong pathogenic variants across 4,780 genes enriched for disease association (1.6 for Admixed American and Middle Eastern, 1.7 for African and East Asian, 1.8 for Central/South Asian and European) and 2.4-2.9 per genome among all genes across ancestries (2.4 for Middle Eastern, 2.5 for Admixed American, 2.8 for European and East Asian, 2.9 for Central/South Asian and African).

This result matches that of a similar analysis conducted among 300 rare-disease probands [45] that did not consider ancestries. Another publication found 4.3 and 4 pathogenic variants per person for European and non-European genomes across all genes, respectively, in the All of Us cohort [9], suggesting that our results align with previous research and that pathogenic variation per genome is largely consistent across ancestries. Previous studies have also demonstrated that selection does not differ among genetic ancestries [11], that (strongly) pathogenic variants are expected to be similarly rare in all ancestries [46], and that deleterious load is resistant to recent population history [41]. Additionally, both recently expanded and non-expanded populations are likely to carry the same deleterious genetic load [17]. While we expect that benign genomic variation will vary considerably between populations due to previous founder events, bottlenecks, and population size differences, the accumulation of pathogenic variants is affected by additional factors, including natural selection. Interestingly, variants classified by the predictors as indeterminate are observed to differ between the populations, from 22.3 variants for Admixed American to 34.9 variants for African ancestry (Figure 2a), which leads to the largest VUS per-person numbers in the individuals of African ancestry.

Finally, we assessed the performance of variant impact prediction tools using ClinVar and HGMD data. We estimated AUC on these datasets stratified by ancestry and allele frequency (Figure 3). Binning by allele frequency is essential, as the extreme sample size differences enrich the European population for much rarer variants that are more likely to be pathogenic (Figure 2) and also more likely to be correctly predicted as pathogenic (Figure 3). This is evident when comparing across bins, where AUC dramatically decreases as the allele frequency increases (Figure 3). We find that few differences exist in performance in the European-ancestry group compared to non-Europeans, with only a few tests being statistically significant and with small effect size. We conclude that these results present evidence in favor of fair performance of computational tools, and suggest that, on the whole, they perform largely the same across ancestral groups. However, the few statistically different results need to be further investigated.

Accurately and fairly performing ancestry-specific evaluation involves several challenges. One limitation is a large sample size discrepancy between European and non-European genomes in the UK Biobank. This is also true for gnomAD, leading to a situation where the allele frequencies for the European population are more accurately estimated from gnomAD than those of non-Europeans, especially for the smallest populations. This presents a potential bias in our analysis, as rarity of variants has a large effect on their potential for pathogenicity. Our choice to bin the analysis by allele frequency removes major confounding, but it cannot control for incorrectly determined frequencies that may exist, affecting the AUCs.

In summary, we find that the performance of computational tools on the prediction of the impact of rare missense variants is similar across six ancestral groups. We also report a comparable number of predicted pathogenic rare missense variants across all genomes, irrespective of ancestry. We conclude that, while imperfect, these tools can be reliably used in clinical variant classification regardless of the genetic ancestry of an affected individual.

## Methods

### Data

#### The UK Biobank resource

The UK Biobank contains a cohort of 502,536 participants from England, Wales, and Scotland recruited between 2006 and 2010, with participant age ranging from 40-69 years at recruitment [1, 47]. DNA was extracted from whole blood and sequenced by the Regeneron Genetics Center (RGC). Exome sequencing data were available for 469,765 participants.

### ClinVar and HGMD data

Missense variants were extracted from ClinVar [26], downloaded on December 08, 2025, and the Human Gene Mutation Database (HGMD) Pro 2025.1 [44].

### Participant definition

Quality control was performed by RGC as previously described [51]. Briefly, individuals with evidence of contamination, unresolved duplications, sex discrepancies, and discordance between exome sequencing and genotyping data were removed. Participants were grouped by Pan-UKB designations [23], and annotated as being of European, African, Central/South Asian, East Asian, Admixed American, or Middle Eastern genetic ancestry. Individuals not categorized by Pan-UKB were not included in this analysis. Additionally, genetic similarity was determined by RGC using PRIMUS [43]. Highly genetically related individuals, including all first- and second-degree relatives, and some third-degree relatives were removed. After filtering, we conducted the analysis with 406,951 European, 8,542 Central/South Asian, 6,334 African, 2,637 East Asian, 1,514 Middle Eastern, and 963 Admixed American genetic ancestry participants. Birth country (UK Biobank data field 20115) was reported for participants who indicated being born outside of the United Kingdom or Republic of Ireland.

### Variant processing and curation

Exome sequencing variant processing was conducted as described previously [20]. Briefly, exome variants with genome quality above 20 and depth above 10 for SNVs were collected across all included participants and annotated using VEP version 114 [30]. Alternate allele balance (AB) was calculated using Picard [40] and variant sites with AB above 0.15 for SNVs were excluded. Per-population filtering included removing variants with Hardy-Weinberg equilibrium P-value above 10^*−*10^ and missingness below 2%. Variant consequences were defined by their effect in the MANE-select transcript [32]. In rare cases, more than one MANE-select transcript exists for a given variant, in which case the variant was defined by its maximum reported order of severity as calculated by Ensembl [19].

Variant allele frequencies were calculated using gnomAD [22] version v4, using exome-derived allele frequencies “afr”, “amr”, “eas”, “mid”, “sas”, and “nfe” for the African, Admixed American, East Asian, Middle Eastern, Central/South Asian, and European genetic ancestry populations, respectively. This analysis was carried out on all genes and also restricted to 4,780 disease genes, defined here as genes with at least one 1-star or higher pathogenic or likely pathogenic variant in ClinVar. All variants were mapped to the GRCh38 reference panel.

### Variant interpretation tools and ACMG/AMP evidence strengths

REVEL [21], MutPred2 [35] and AlphaMissense [7] were selected as the primary variant effect predictors for this study, while VEST4 [6], BayesDel (without allele frequency) [15], VARITY [53], and ESM1b [4] were selected for the secondary analysis. The selection was based on their performance in the Critical Assessment of Genome Interpretation [50, 38], availability of ClinGen-recommended calibration thresholds [34, 3], methodological diversity and our ability to exclude training data from the evaluation dataset. All predictor scores were returned based on the score reported in the MANE-select transcript [32], or canonical transcript if a MANE-select was not available. For each method, missense variants were stratified according to the evidence strengths defined in the ACMG/AMP recommendations based on the score intervals for mapping raw prediction scores into discrete evidence strengths [34, 3]. With this step, variants were assigned direction (benign, pathogenic) and strength (supporting, moderate, strong, very strong) of evidence. Those that were not assigned a direction-strength pair were designated as indeterminate. For compatibility with the upcoming ACMG/AMP guidelines, evidence was also presented using the point-based system, with supporting evidence being assigned ±1 point, moderate evidence split into ±2 and ±3 points, and strong evidence presented as ±4 points. The sign in the point system reflects the direction of evidence (+ for pathogenic; *−* for benign).

### Evolutionary constraint computation

We selected GERP++ rank scores [8], developed over multiple mammalian species, as our primary measure of evolutionary constraint. PhyloP [36], another measure of evolutionary constraint over multiple vertebrate species, was used as a secondary measure. Approximately 0.7% of missense variants lacked GERP++ (PhyloP = 0.03%) scores and were removed from all analyses where the tool was applied.

### Performance evaluation

To accurately assess the performance of predictors on labeled ClinVar and HGMD variants, we removed any data that were used in model training. When assessing REVEL, MutPred2, and BayesDel, we removed 7,345, 1,502, and 10,844 variants used in their training sets, respectively. AlphaMissense was not trained with direct supervision using pathogenic and benign variants, and hence no variants were removed when assessing performance.

The performance of predictors in discriminating pathogenic from benign variants was evaluated by calculating the area under the ROC curve (AUC), where the positive class label was assigned to variants classified as “pathogenic,” “pathogenic/likely pathogenic,” “likely pathogenic,” and “DM,” and the negative class label was assigned to variants classified as “benign,” “benign/likely benign,” and “likely benign.” In 461 instances where HGMD and ClinVar provided conflicting class labels, the ClinVar designation was retained.

### Ethics statement

The UK Biobank study was approved by the National Health Service National Research Ethics Service and all participants provided written informed consent to participate in the study for health-related research. Information about ethics oversight in the UKB study can be found at https://www.ukbiobank.ac.uk/ethics/. This study has been conducted using the UK Biobank resource application 26041.

### Statistical computing

Data management and analytics were performed using the REVEAL/SciDB translational analytics platform from Paradigm4.

## Acknowledgments

We thank Drs. Joseph Rothstein and Weiva Sieh for filtering out variants in our set that were present in REVEL’s and constituent tools’ training sets. We thank Dr. Bing-Jian Feng for sending us the set of variants used to train BayesDel.

